# MultiEditR: An easy validation method for detecting and quantifying RNA editing from Sanger sequencing

**DOI:** 10.1101/633685

**Authors:** Mitchell Kluesner, Annette Arnold, Taga Lerner, Rafail Nikolaos Tasakis, Sandra Wüst, Marco Binder, Branden S. Moriarity, Riccardo Pecori

## Abstract

RNA editing is the base change that results from RNA deamination by two predominant classes of deaminases; the APOBEC family and the ADAR family. Respectively, deamination of nucleobases by these enzymes are responsible for endogenous editing of cytosine to uracil (C-to-U) and adenosine to inosine (A-to-I). RNA editing is known to play an essential role both in maintaining normal cellular function, as well as altered cellular physiology during oncogenesis and tumour progression. Analysis of RNA editing in these important processes, largely relies on RNA-seq technology for the detection and quantification of RNA editing sites. Despite the power of these technologies, multiple sources of error in detecting and measuring base editing still exist, therefore additional validation and quantification of editing through Sanger sequencing is still required for confirmation of editing. Depending on the number of RNA editing sites that are of interest, this validation step can be both expensive and time-consuming. To address this need we developed the tool MultiEditR which provides a simple, and cost-effective method of detecting and quantifying RNA editing form Sanger sequencing. We expect that MultiEditR will foster further discoveries in this rapidly expanding field.

## INTRODUCTION

The genetic information contained in DNA defines, via RNA translation, the amino acid composition of a protein. Epitranscriptomics as a field studies modifications in RNA leading to changes in features of a transcript, such as translational efficiency, stability, localization, alternative splicing choice and amino acid composition.

RNA editing is defined as a change in nucleotide content within the RNA molecule. Editing was initially coined as a term to describe a situation in *Trypanosoma brucei* where insertion of uridines within mRNA helped define the reading frame of a transcript (Benne et al. 1986). In contrast, RNA editing in mammals predominantly refers to a base change that is the result of RNA deamination by two families of enzymes: the APOBEC (Apolipoprotein B mRNA Editing Catalytic Polypeptide-like) protein family and the ADAR (Adenosine Deaminase that Acts on RNA) protein family.

APOBEC deaminases are responsible for cytosine to uracil (C-to-U) editing (Wedekind et al. 2003). C-to-U RNA editing was initially discovered in intestine of different mammals (Chen et al. 1987; Powell et al. 1987) as a consequence of the ability of the APOBEC1 protein to recode within a specific transcript (the *Apob* transcript) a CAA codon (Q2152 in humans) to a stop codon, resulting in a truncated form of the APOB protein (APOB-48) (Navaratnam et al. 1993; Teng, Burant, and Davidson 1993). This shorter version of APOB together with APOB-100 (the long, unedited form) are essential for lipid metabolism: the former is the main protein component of chylomicrons and the latter, the main component of low density lipoproteins (LDLs) (Yao and McLeod 1994; Young 1990).

The RNA editing activity of APOBEC1 is not limited to *Apob* mRNA. In fact APOBEC1 edits many more transcripts in both *Apob* expressing tissues and tissues where *Apob* is not present, leading to C-to-U RNA editing in hundreds of transcripts mainly in their 3’UTR (Blanc et al. 2014; Harjanto et al. 2016; Rayon-Estrada et al. 2017; Rosenberg et al. 2011). APOBEC1 C-to-U editing seems to have a role also in maintaining brain physiology in mice, rats and humans (Cole et al. 2017; Kankowski et al. 2018; Meier et al. 2005), suggesting a critical role of C-to-U editing in the regulatory mechanism for brain homeostasis (Gagnidze et al. 2018). Finally, human APOBEC3A and APOBEC3G were also shown to be able to perform C-to-U RNA editing (Sharma et al. 2015, 2016, 2017, 2019), suggesting that in humans, three APOBECs are able to catalyze this epitranscriptomic mark.

ADAR proteins (ADAR1 and ADAR2 in humans) are responsible for adenosine to inosine (A-to-I) editing. Inosines are interpreted by the cells as guanosines (G) (Lengyel, Speyer, and Ochoa 1961) resulting in an A-to-G base change. A-to-I editing can be divided in “recoding” editing occurring in protein-coding region and “not-recoding editing” in the non-coding part of the transcriptome. In humans, more than a thousand recoding sites have been documented (Picardi et al. 2017; Ramaswami and Li 2014). This kind of editing is performed by mainly ADAR2 and is principally present in neural tissues where editing of ion channels and neuroreceptors is essential for physiological functioning of the brain (Rosenthal and Seeburg 2012). However, the vast majority of editing does not affect recoding. Rather, it can alters splicing (Lev-Maor et al. 2007), miRNA binding sites in 3’UTR region (Pinto et al. 2018) or the cognate miRNAs themselves (Kawahara et al. 2007). Most importantly, this editing is required to prevent activation of cytosolic immune innate system against endogenous dsRNA, representing, the main function of ADAR1 editing (Liddicoat et al. 2015; Mannion et al. 2014; Pestal et al. 2015).

Overall, physiologic RNA editing plays an essential role in neuronal and immune processes. Indeed knock out mice for ADAR1 or ADAR2 have lethal phenotypes (Hartner et al. 2004; Wang et al. 2004; Ward et al. 2010) and knock out mice for APOBEC1 exhibit myelin fragmentation and lysosome/autophagosome accumulation in microglia and neurons, leading to an early-aging phenotype characterized by cognitive and motor deficits (Cole et al. 2017). Additionally, A-to-I RNA editing is the most abundant modification in mRNA (Picardi et al. 2017; Ramaswami and Li 2014) both in healthy tissues and during oncogenesis and tumour progression (Fumagalli et al. 2015; Galeano et al. 2012; Han et al. 2015; Lin and Chen 2019; Paz-Yaacov et al. 2015; Xu and Öhman 2019).

Furthermore, there are emerging therapeutic contexts of this modification. Specifically, site-directed (or targeted) RNA editing is a prospective method for transient reprogramming of genetic information as a potential alternative to permanent genomic DNA editing (Vogel and Stafforst 2018). RNA editing is also substantially more efficient than classical CRISPR/Cas9 techniques and consequently, many labs have recently invested in developing targeted editing technologies (F, Montiel-Gonzalez et al. 2013; F, Montiel-González, C, Vallecillo-Viejo, and Rosenthal 2016; Merkle et al. 2019; Monteleone et al. 2019; Stafforst and Schneider 2012; Vallecillo-Viejo et al. 2018; Vogel et al. 2018; Wettengel et al. 2017). Perhaps the most promising of these therapeutic approaches is to redirect endogenous ADARs using various CRISPR enzymes and guide RNAs to specific sites, such as for “mutation restoration” events (RESTORE; (Merkle et al. 2019)). Overall, therapeutic application of RNA editing appears to be a promising avenue.

Current identification of RNA editing is mediated by RNA-seq in tandem with several different algorithmic approaches (Eisenberg and Levanon 2018; Ramaswami and Li 2016). These approaches largely rely on the comparison of DNA-to-RNA or RNA-to-RNA sequences in which editing is detected as C-to-T or A-to-G base conversions. In the case of RNA-to-RNA comparisons, RNA from a WT sample is compared to RNA from a sample containing a KO for an RNA editing enzyme of interest. In theory this approach is straightforward, however in practice there are many sources of base mismatches in NGS data (Eisenberg and Levanon 2018). Therefore validation and quantification of editing through Sanger sequencing and bacterial colony sequencing of subcloned polymerase chain reaction (PCR) amplicons are essential before “true” editing sites are defined. Depending on the number of RNA editing sites that are of interest, this validation step can be both expensive and time-consuming. Consequently, it would be of use to this rapidly expanding field if a tool existed to efficiently, and cost effectively, detect and quantify RNA editing.

Here we have developed, and validated a software tool (MultiEditR) that provides simple, and cost-effective quantification of RNA editing levels by accurately “calling” editing sites from chromatograms (from standard fluorescence-based Sanger sequencing of amplicons). Use of this program cuts down time and cost, increasing the number of validations that can be done. Considering the importance of these methods and their probable future use in therapeutics we expect MultiEditR to become a very useful tool for the RNA editing community, initially, but also the RNA modification community more broadly.

## METHODS

### Plasmids

The mCherry-mApob-eGFP plasmid (CmAG) was obtained by substituting the human *APOB* with mouse *Apob* in the original plasmid mCherry-Apob-eGFP (Severi and Conticello 2015). This plasmid (kind gifts from Dr. Silvestro Conticello, Florence, Italy) was digested with HindIII-SmaI and a PCR fragment of mouse *Apob* (467bp from RNA of jejunal epithelial cells from the small intestines of C57BL/6 (Rosenberg et al. 2011), oligos #1-2) was inserted into the plasmid using NEBuilder® HiFi DNA Assembly Master Mix (NEB).

The mouse APOBEC1 expression vector (pCMV APOBEC1) was a kind gift of Dr. Dewi Harjanto (Laboratory of Lymphocyte Biology, The Rockefeller University).

The mouse RBM47 expression vector (pCMV RBM47) was obtained by inserting a PCR fragment containing the coding sequence of mouse RBM47 (transcript variant 4, mRNA Sequence ID: NM_001291226.1) into the mCherry-Apob-eGFP cut with NheI-BsrGI. The amplification was done using oligos #3-4 on RNA of jejunal epithelial cells from the small intestines of C57BL/6(Rosenberg et al. 2011)and the cloning with NEBuilder® HiFi DNA Assembly Master Mix (NEB).

LentiCRISPRv2 was a gift from Feng Zhang (Addgene, plasmid #52961; http://n2t.net/addgene:52961; RRID:Addgene_52961)(Sanjana, Shalem, and Zhang 2014)DNA oligos #5-6 and #7-8 were cloned into this plasmid following the “lentiCRISPRv2 and lentiGuide oligo cloning protocol” (Addgene plasmid #52961) to generate lenti-CRISPR-ADAR1 exon 3 and exon 2, respectively. As a non-editing transduction control, lenti-CRISPR-NT (Lenti-NT) was cloned accordingly, using oligos #9-10.

pCMV-DR8.91 (coding for HIV gag-pol) and pMD2.G (encoding the VSV-G glycoprotein) were kind gifts from Prof. Didier Trono, Lausanne, Switzerland.

pSpCas9(BB)-2A-GFP (PX458) was a gift from Feng Zhang (Addgene plasmid # 48138; http://n2t.net/addgene:48138; RRID:Addgene_48138). The plasmid was digested with BsbI (NEB) and dephosphorylated with RAPID DNA Dephos and Ligation kit (Roche). Oligos #11-14 are all 5’ phosphorylated. The oligo pairs #11-12 and #13-14 containing complementary sequences were annealed to each other then ligated to the dephosphorylated PX458 to generate plasmids PX458-iv-sgRNA-A1 11 (cutting in exon 4) and PX458-iv-sgRNA-A1 39 (cutting in exon 5) respectively.

### Cell lines and transfections

A549 cells (A-549, RRID:CVCL_0023, DKFZ Germany) were cultured at 37°C, 5% CO_2_, in high glucose DMEM (Sigma), supplemented with 10% fetal bovine serum (FBS, PAN Biotech) and Penicillin/Streptomycin (Sigma).

HEK293T cells (ATCC-CRL-3216) were cultured at 37°C, 5% CO_2_, in high glucose DMEM (Sigma), supplemented with 5% fetal bovine serum (FBS, PAN Biotech) and Penicillin/Streptomycin (Sigma).

RAW 264.7 cells (ATCC® TIB-71™) were subcultured at 37°C, 5% CO_2_, in high glucose DMEM (Sigma-Aldrich), supplemented with 5% endotoxin low fetal bovine serum (Sera Pro FBS, PAN Biotech), 1% Glutamine and 1% Penicillin/Streptomycin (Sigma).

The cell lines were regularly tested for mycoplasma contamination in our facility Multiplexion (F020, DKFZ) (www.multiplexion.de)

### Generation of A549 ADAR1 knock out cell line

Lenti-CRISPR-ADAR1 exon 3, exon 2 or NT in combination with pCMV-DR8.91 and pMD2.G were calcium-phosphate transfected in HEK293T cells for lentiviral particle production (ratio 3:1:3). 48-72h post transfection cell-free supernatant was harvested and used for transduction of A549 cells. The transduced cells were selected with puromycin (1 μg/ml). Immediately after the selection control (non-transduced A549) died, limiting dilution in 96-well plates was performed for ADAR1 KOs (0.5 cells/well) and clonality was validated by visual inspection with a microscope; the Lenti-NT control was kept polyclonal. KO of ADAR1 was validated by Western blot (Anti-Human ADAR1 [D7E2M] rabbit mAb, Cell Signaling Technology, cat. no.#14175). Two clones, number 5 and 7, resulted in completely abolished ADAR1 (p110 and p150) expression (Figure S1). For further experiments we used only clone #5.

### Generation of RAW 264.7 APOBEC1 knock out cell line

PX458-iv-sgRNA-A1_11 and PX458-iv-sgRNA-A1_39 were cotransfected using Amaxa® Cell Line Nucleofector® Kit V (Lonza) into RAW 264.7 cells following the manufacturer’s protocol for RAW 264.7 cells and a Nucleofector™ 2b Device (Lonza). 48h post transfection GFP positive cells were single cell sorted into 96 well plates and clonality was validated by visual inspection with a microscope. Clones were screened by amplifying targeted regions from genomic DNA (produced by High Pure PCR template preparation kit (roche)) using oligos #15-16 and #17-18 then sanger sequencing. This was followed by additional cloning of amplified region using CloneJET PCR Cloning Kit according to the manufacturer’s instructions and transforming DH5α bacteria with ligated product. 10 resultant bacteria colonies were sent for sequencing to determine genetic changes to targeted region. One clone which was subsequently used contained in the region targeted by PX458-iv-sgRNA-A1_39 either a single base pair deletion or a two base pair deletion. Knockout was further confirmed by RT PCR (using Onestep RT-PCR kit – Qiagen) amplification of B2m 3’UTR region from extracted RNA defined by oligos #19-20 known to be edited and determining absence of editing compared to amplified region from the parental cells (Figure S2).

### RNA extraction, DNAse treatment and RT-PCR

RNA was extracted using RNeasy Mini kit (Qiagen), and treated with DNase (Turbo DNA-free kit, Invitrogen). All the PCRs on RNA were performed with gene specific primers (listed in Table S1) and OneStep RT–PCR kit (Qiagen).

### Titration experiment

For the C-to-U editing HEK293T were transfected with CmAG(50ng), APOBEC1(200ng) and RBM47(200ng) expression vectors or CmAG(50ng) only. For transfection we used a mix of plasmidic DNA and Polyethylenimine (PEI) in a ~1:4 ratio (450ng of DNA: 2μg PEI). 72h after transfection we extracted RNA and amplified *Apob* (oligos #21-2). This allowed us to obtain *Apob* fragments heavily edited or not edited, respectively. These two fragments were cloned into CloneJET PCR Cloning Kit (Thermo Scientific) and several colonies were screened by sequencing. From this screening we obtained two pJET vector containing *Apob* with no editing (pJET-CmAG-WT) and 6 edited sites (pJET-CmAG-6x). These two vectors were then mixed together in titrated amounts from 0% to 100% and subjected to capillary Sanger sequencing with universal primers pJET1.2 forward and reverse.

### RNA-seq

We prepared RNA-seq libraries in duplicate from A549 wild type and *ADAR1* knock out clone 5 (Figure S1), and in triplicates from RAW 264.7 wild type and RAW 264.7 APOBEC1 knock out.

Total RNA was extracted from 10,000,000 cells in duplicate (A549 wt and ADAR1 ko) or triplicate (RAW 264.7 and RAW 264.7 APOBEC1 knockout each from separate plates). RNA was extracted using RNeasy mini kit (Qiagen) and then treated with turbo DNase (Life technologies). RNA concentration and integrity was determined by a Qubit 4 (Thermo Fischer scientific) using the Qubit RNA BR assay kit or Qubit XR assay kit and the Qubit RNA IQ kit (Thermo Fischer scientific). 1μg of RNA was processed with Kapa mRNA hyperprep Kit for illumine platforms (Kapa Biosystems, Roche) and KAPA Single-Indexed Adapter Kit for Illumina® Platforms (Kapa Biosystems, Roche).

Libraries were sequenced with the Illumina HiSeq 2000 v4 technology, generating 125-nucleotide, paired-end reads. Adapters were trimmed using the Trim Galore software (https://github.com/FelixKrueger/TrimGalore). Before and after trimming we evaluated the RNAseq quality with FastQC (https://www.bioinformatics.babraham.ac.uk/projects/fastqc/). Quality control, including per-base quality, duplication levels, and over-representative sequences, passed all the checkpoints. The RNA-seq reads were then aligned to hg19 (A549 data) or mm10 (RAW 264.7 data) reference genomes (publicly available by the UCSC genome browser) using the STAR aligner (v. 2.6.0a, https://github.com/alexdobin/STAR(Dobin et al. 2013), with default settings. Potential PCR duplicates were removed using the MarkDuplicates function from Picard tools (http://broadinstitute.github.io/picard).

The aligned RNAseq (.bam files) were sorted and indexed with SAMtools (https://github.com/samtools/samtools). The sorted bam files were used as an input for the REDItoolsDnaRna.py script, part of the REDItools suite (v. 1.0.4(Picardi et al. 2015; Picardi and Pesole 2013). REDItoolsDnaRna.py performs a comparative position-per-position analysis in parallel between an RNA and a DNA bam files, so as to eliminate variants on the RNA, the signal of which derives from the genomic DNA. For this analysis however, we employed RNA-seq from A549 ADAR1 wild type and ADAR1 knock out, or RAW 264.7 wild type and RAW 264.7 APOBEC1 knock out. Sequences from knock out were used as background, in order to identify the editing events for which ADAR1 or APOBEC1 is responsible for. The options we set for a genomic position to be considered for variant calling, required minimum coverage of 5 reads, with at least 3 reads supporting the editing event, minimum FastQ offset value 33, per-base. The aforementioned settings were used to run the analysis per pair and specifying precise gene’s coordinates with the option -Y. Genes and their coordinates are listed in the Supplementary information together with output file from REDitool analysis.

### MultiEditR development

MultiEditR was written in the statistical programming language R (v. 3.4.2) using RStudio (v. 1.1.383). The MultiEditR web app was developed using R shiny (https://shiny.rstudio.com/). global.R contains the main runEditR() function which takes input files and parameters and returns the output analysis, dependencies.R specifies the required packages for the web app, most notably *sangerseqR* for reading and analyzing .ab1 files (https://bioconductor.org/packages/release/bioc/html/sangerseqR.html), *gamlss* for zero adjusted gamma distribution modelling (https://www.gamlss.com/), *tidyverse* packages for data manipulation and visualization (https://www.tidyverse.org/), and *shiny* for support of the web application. helpers.R contains functions for processing the data throughout the analysis which are called by global.R. server.R interfaces the inputs from the user side to the server side to be ran through runEditR(), and ultimately returns outputs to the user visual interface. ui.R specifies the visual interface of the web app. All code is available at https://github.com/MoriarityLab/MultiEditR (See Figure 2A for a visual layout of program).

### Data analysis

All statistical analyses were performed in R studio. The level of significance was set at α = 0.01. Student’s one-sample, two-tailed t-tests were used as indicated in the text. Data were visualized in R studio employing various tidyverse (https://www.tidyverse.org/) and Bioconductor (https://www.bioconductor.org/) packages. See code at https://github.com/MoriarityLab/MultiEditR for reproducible analysis.

## RESULTS

### Titration analysis of multiple edits

Previously we developed Edit deconvolution by inference of traces in R (EditR) for the quantification of base conversions within a single, discrete region of the chromatogram. However, we wanted to assess whether a similar approach was applicable to the detection and quantification of multiple conversions from endogenous RNA editing. To determine if RNA editing can be accurately quantified from Sanger sequencing we extracted RNA from 293T cells after transfection of CmAG, APOBEC1 and RBM47 expression vectors or CmAG only. From these samples we amplified the Apob region of the CmAG transcript, which is established to be highly edited(Yamanaka et al. 1996) (Figure S3). Blunt end cloning of the PCR product yielded a plasmid with six C-to-T mutations (pJET-CmAG-6x) and a plasmid without any edits (pJET-CmAG-WT). pJET-CmAG-6x was titrated with pJET-CmAG-WT from 0% to 100% in increments of 5%, with smaller 2.5% increments at the low- and high-end of the titration. Titrations were subjected to Sanger sequencing and base conversion was measured by calculating the percent height of each base (Figure 1A-D). Both C>T and G>A titrations yielded well fit linear regressions of Sanger observed vs. expected values (R^2^_C_ = 0.981, R^2^_T_ = 0.981, R^2^_G_ = 0.983, R^2^_A_ = 0.983). Furthermore, measuring the percent height of each base was able to discriminate base conversions as low as 7.5% from background (Figure 1B, 1D), consistent with previous work (Kluesner et al. 2018).

**Figure 1.**
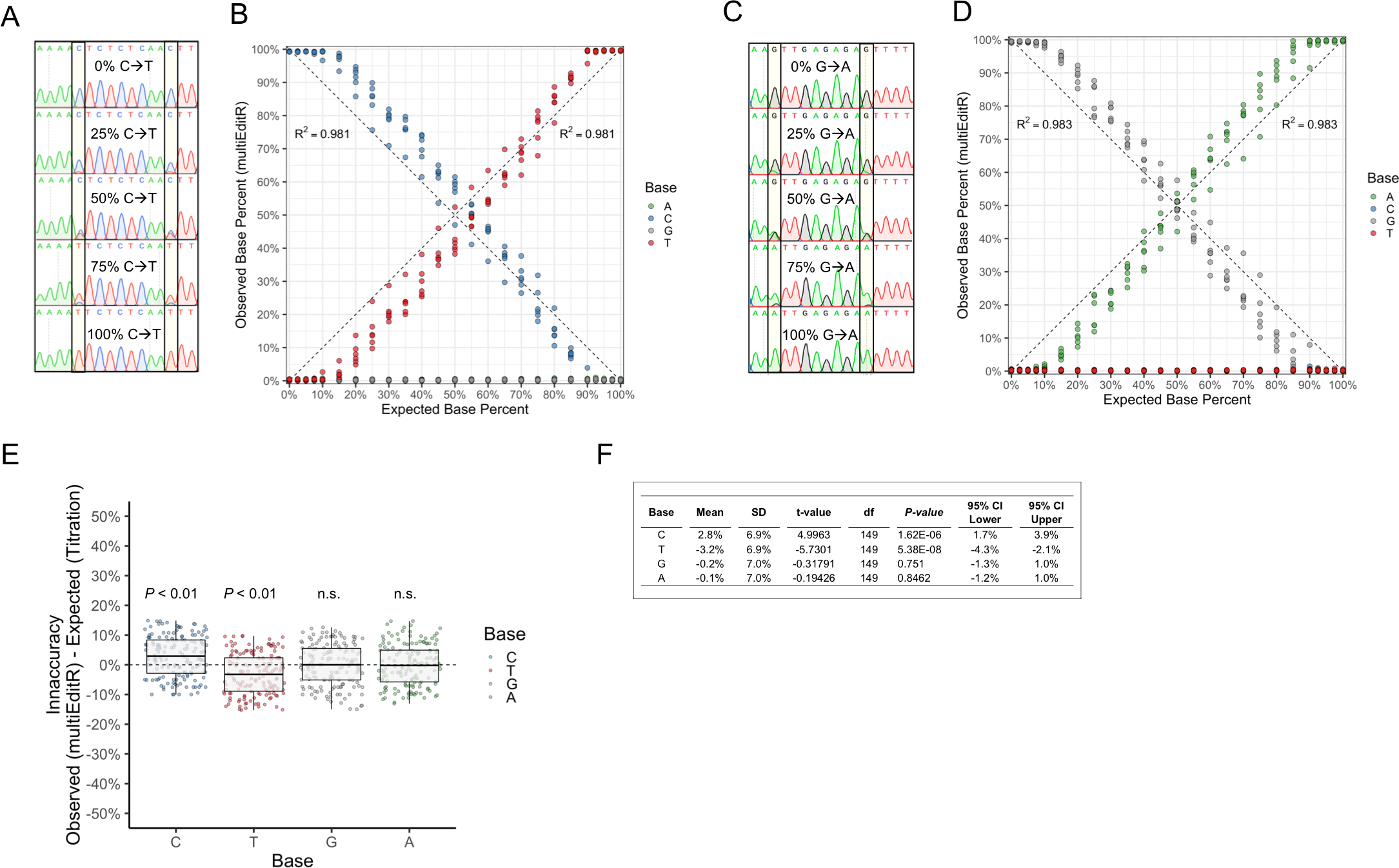
Multiple base conversions can be quantified from Sanger sequencing. **(A)** Chromatograms from C-to-T titration showing a change in peak height at two sites. **(B)** Titration of pJET-CmAG-WT with pJET-CmAG-6x sequenced with the reverse primer, coefficient of determination was calculated relative to the identity line, N = 6 sites per chromatogram. **(C)** Chromatograms from G-to-A titration showing a change in peak height at two sites. **(D)** Titration of pJET-CmAG-WT with pJET-CmAG-6x sequenced with the forward primer, coefficient of determination was calculated relative to the identity line y = x, N = 6 sites per chromatogram. **(E)** Inaccuracy of MultiEditR relative to expected titration values, significance was determined using Student’s one-sample *t*-test relative to an inaccuracy of 0%. **(F)** Table of results from Student’s one-sample *t*-test of inaccuracies.

**Figure 2.**
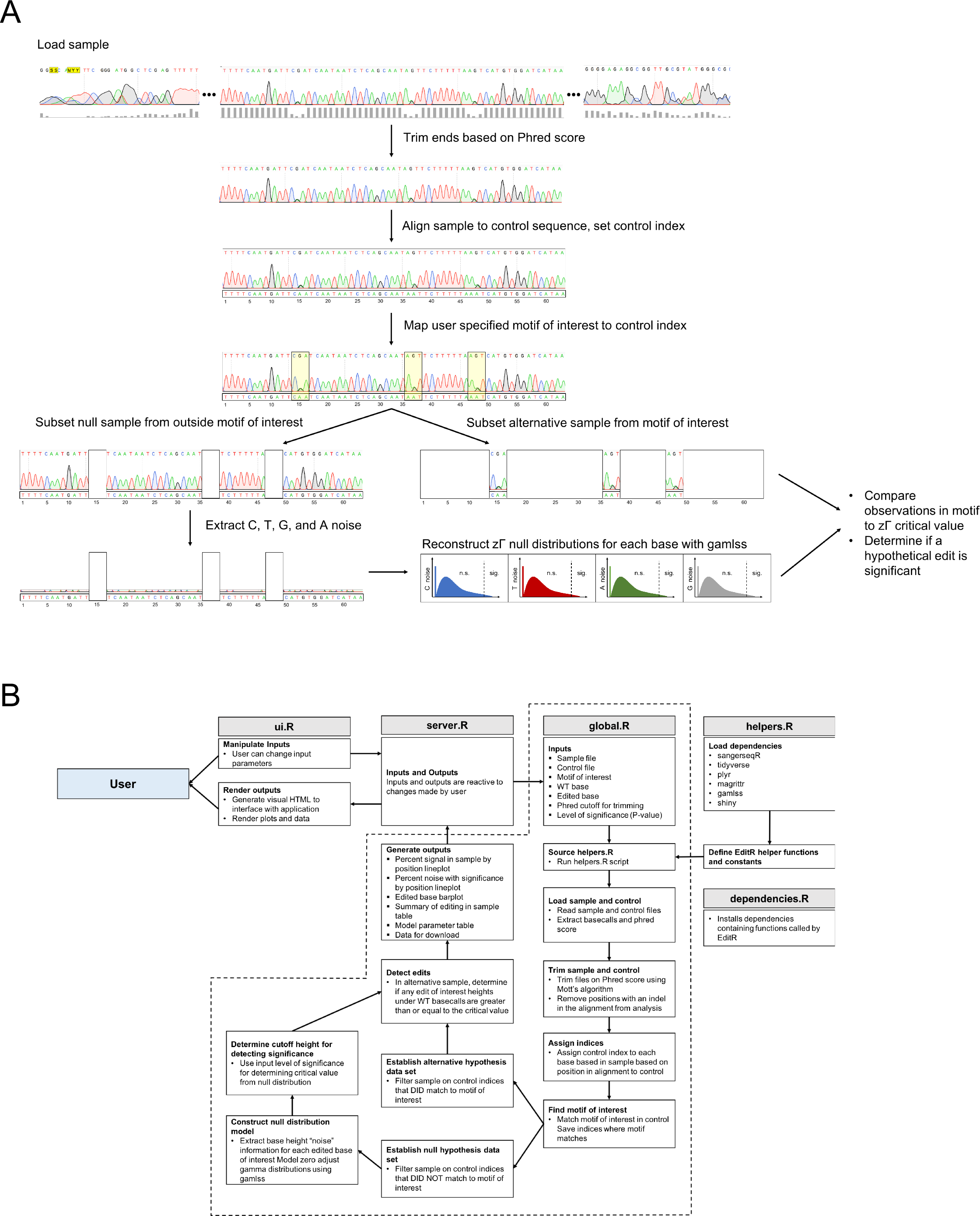
MultiEditR algorithm. **(A)** Visual algorithm with showing zero adjusted gamma (zΓ) distribution generation, and null hypothesis significance testing approach. Algorithm can be applied to any motif(s), WT base(s) and edited base(s). **(B)** MultiEditR algorithm flow chart in the context of the MultiEditR web app made using R Shiny (https://github.com/MoriarityLab/MultiEditR).

Accuracy was further assessed by calculating the difference in observed and expected values for each base, i.e. inaccuracy, and performing Student’s one-sample *t*-test (Figure 1E). The mean inaccuracy of G and A bases was not significantly different from 0% (*P*_*G*_ = 0.751, *P*_*A*_ = 0.8462), meanwhile the mean inaccuracy of C and T bases was significantly different (*P*_*C*_ = 1.62 × 10^−6^, *P*_*T*_ = 5.38 × 10^−8^), with C overestimating and T underestimating. Although C and T bases were significantly different, the magnitude of this difference was less than 5% (M_C_ ± SD = 2.8% ± 6.9%, M_T_ ± SD = −3.2% ± 6.9%). Based on these results indicating that measuring percent base height could be used as an accurate method of quantifying RNA editing, we sought to develop a program to automate the process of RNA editing detection and quantification from Sanger sequencing.

### MultiEditR algorithm

Our previous work on EditR to detect and quantify single-site CRISPR-Cas9 genome base editing used a null-hypothesis significance testing (NHST) approach, which we hypothesized could be modified to detect and quantify endogenous RNA editing at multiple sites in a single transcript. To address this, we adapted our NHST algorithm from EditR for multiple base conversions spread across an amplicon and called it <u>Mult</u>iple <u>Edit</u> deconvolutions by inference of traces in <u>R</u> (MultiEditR). MultiEditR requires 1) a sample .ab1 file of an amplicon of interest between approximately 300 – 750bp, 2) either a control .fasta file of the sequence of the amplicon of interest without any modifications, or a control .ab1 of the amplicon of interest without any modifications, and 3) a motif of interest consisting of any length of IUPAC nucleotides (e.g. YAR, TCA, NAN, N_20_, N_n_, etc.), 4) a discrete base of interest hypothesized to be edited (e.g. A, C, T, or G), and 5) any hypothesized edited outcomes separated by “|” (e.g. G, T|G, A|T|G, etc.) (Figure 2A).

The MultiEditR algorithm (Figure 2B) begins by loading the sample .ab1 and trimming the ends of the sequence based on a Phred score cutoff (default of 0.0001) using a modified Mott’s algorithm (http://www.phrap.org/phredphrap/phred.html). If a control .ab1 file is used, the file is trimmed in the same manner. Once trimmed, the basecalls are extracted from the sample chromatogram and aligned to the non-edited control sequence, where the control sequence index is joined to the sample. Any positions with indels in the alignment are filtered out of the analysis. The motif of interest is then matched to the control sequence and the control indices where matches are found are used to separate the sample into the alternative sample where matches are found, and the null sample where matches are not found. The noise height of the edited bases of interest (e.g. G if interested in A-to-G edits, or T, G or A if interested in C-to-T|G|A edits) is extracted from the trace of the null sample. These noise samples are used to model zero-adjusted (zΓ) gamma distributions which are used as null distributions for the NHST, wherein the *P*-value determines the critical value within the distribution of calling significance vs. non-significance as previously described (Kluesner et al. 2018). The height of hypothetical edits within the motifs of interests (e.g. height of G under A peaks) are then compared to the critical value generated by the null zΓ distributions. If a hypothetical edit is at or above the critical value it will be called as significant, and reported as an edit. The MultiEditR web application (https://moriaritylab.shinyapps.io/MultiEditR live upon submission, desktop version available at https://github.com/MoriarityLab/MultiEditR), provides diagnostics plots of the sample, visualization and tabulation of detected edits, a summary of the zΓ modelling, and the ability to download the output analysis data as a tab delimited file.

### Comparison of MultiEditR to NGS RNA-seq

After determining that multiple edits can be accurately quantified in a synthetic system, and developing an algorithm for detecting edits, we wanted to validate MultiEditR in the context of endogenous ADAR and APOBEC RNA editing. To test this we performed RNA-seq on the adenocarcinoma cell line A549 and the murine macrophage cell line RAW 264.7, alongside a A549 ADAR1 knockout (KO) cell line (Supplementary Figure 1), and a RAW 264.7 APOBEC1 KO cell line (Supplementary Figure 2). RNA edited sites were identified using the RNA editing analysis tool REDitools (Picardi et al. 2015; Picardi and Pesole 2013) using the KO cell lines as a control. From this, we focused our comparison on four transcripts across the two cell lines that showed ADAR or APOBEC editing. Due to the multiple comparisons problem, first we wanted to assess how well our significance testing approach performed by comparing the observed false positive rate, to that expected by our initial α = 0.01 level of significance. We found that when α = 0.01, our observed false positive rate was substantially higher at 0.29, however the observed false positive rate could be brought below 0.10, and still provide a reasonable sample size (N = 181) when the level of significance was set at α = 0.0001 (Figure 3A).

**Figure 3.**
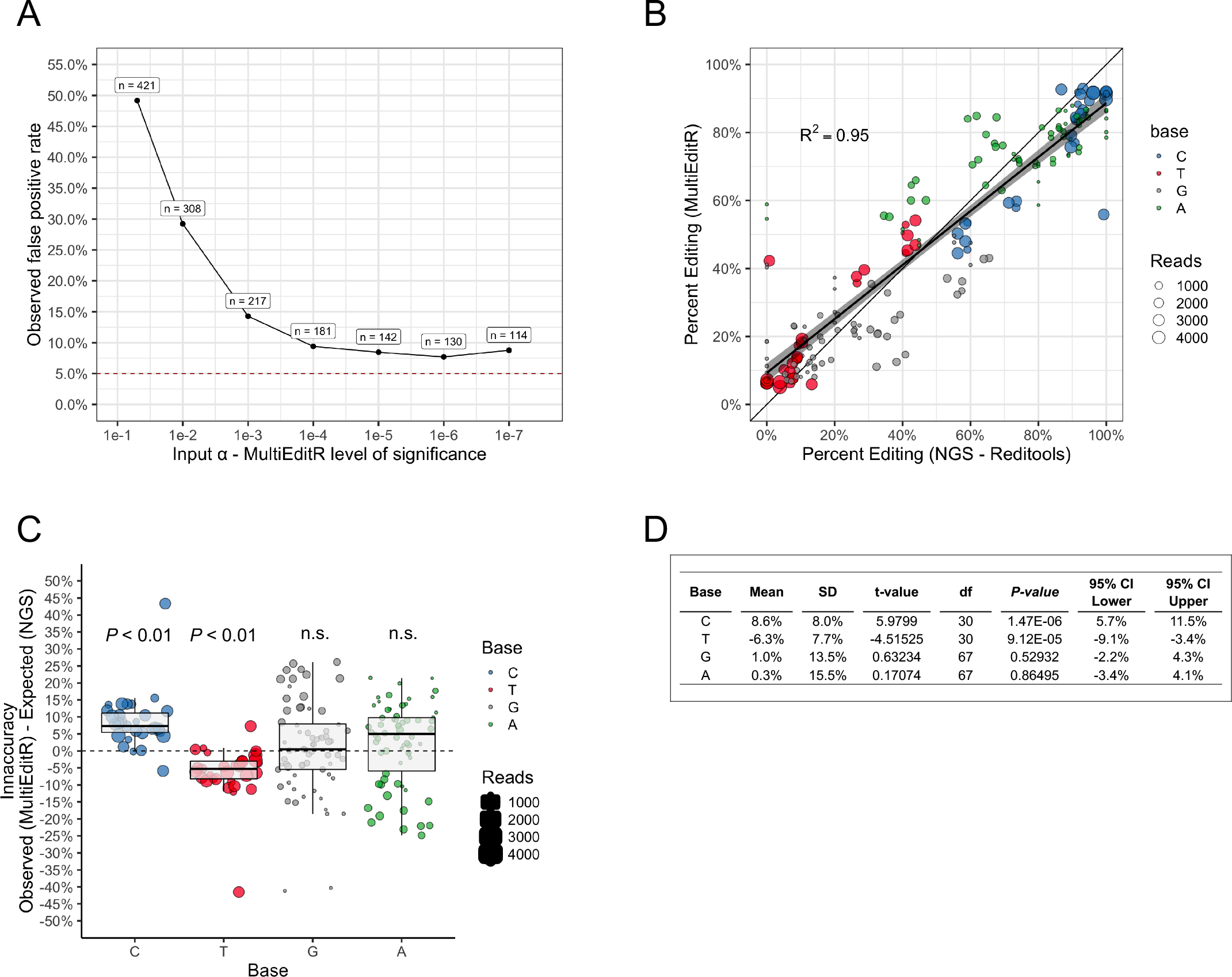
MultiEditR can reliably predict RNA editing relative to RNA-seq. **(A)** False positive analysis of MultiEditR. Line plot shows the observed false positive rate of MultiEditR as a function of the input level of significance (α), i.e. the P-value cutoff. An input level of significance of 0.0001 is required to achieve an observed false positive rate below 0.10. **(B)** Comparison of MultiEditR to NGS RNA-seq analyzed by REDItools. Scatter plot shows percent editing measured by MultiEditR as a function of percent editing measured by REDItools. Coefficient of determination is calculated relative to identity line y = x. Overlaid regression line is a linear regression line-of-best-fit, shaded regions represent 95% confidence interval of the mean. **(C)** Inaccuracy of MultiEditR relative to NGS RNA-seq analyzed by REDItools, significance was determined using Student’s one-sample *t*-test relative to an inaccuracy of 0%. **(D)** Table of results from Student’s one-sample *t*-test of inaccuracies.

In assessing the accuracy of MultIEditR, we found that measurements of editing by MultiEditR compared to REDitools yielded a well fit regression (R^2^ = 0.95) against the identity line (Figure 3B). Accuracy was further assessed by calculating the difference in observed and expected values for each base, i.e. inaccuracy, and performing Student’s one-sample *t*-test (Figure 3C-D). The mean inaccuracy of G and A bases was not significantly different from 0% (*P*_*G*_ = 0.529, *P*_*A*_ = 0.865), meanwhile the mean inaccuracy of C and T bases was significantly different (*P*_*C*_ = 1.47 × 10^−6^, *P*_*T*_ = 9.12 × 10^−5^), with C underestimating and T overestimating relative to REDitools. Although C and T bases were significantly different, the magnitude of this difference was less than 10% but rather tightly dispersed (M_C_ ± SD = −8.6% ± 8.0%, M_C_ ± SD = 6.3% ± 7.7%), suggesting a systematic error in measuring these bases. In contrast, although G and A bases were not significantly different from REDitools measurements, the standard deviation of inaccuracy was quite large (M_G_ ± SD = −1.0% ± 13.5%, M_A_ ± SD = −0.3% ± 15.5%) suggesting a greater influence of random error in these measurements. Collectively, these results indicate that MultiEditR can robustly detect endogenous RNA editing, while producing reasonably precise measurements of editing, especially when measuring C-to-T editing (or A-to-I, antisense T-to-C). However, further work is needed in assessing the magnitude and source of base specific error when comparing MultiEditR to REDitools analysis of RNA-seq.

## DISCUSSION

RNA editing plays an important role in physiological and pathological processes. For this reason detection, validation and quantification of editing are essential. Improvements in RNA-seq technology have allowed the detection of a vast amount of editing sites (Blanc et al. 2014; Picardi et al. 2017; Ramaswami and Li 2014; Rosenberg et al. 2011) which are identified from RNA-seq data as C-to-T or A-to-G base mismatches relative to a reference sequence. Considering the many sources of mismatches (Eisenberg and Levanon 2018) and the needs of bacterial colony sequencing of PCR amplicons for precise quantification, depending on the number of RNA editing sites to validate this step can be both expensive and time-consuming.

To overcome these limitations we first assessed the ability to use percent base height as an accurate method of quantifying RNA editing of several bases in the amplicon. Our results show a well fit linear regression of Sanger observed vs. expected values both for C>T and G>A base changes, being able to discriminate base conversions as low as 7.5% from background (Figure 1B, 1D), consistent with previous work (Kluesner et al. 2018). We also compared the accuracy of the measured editing result. We also found that editing for G and A bases, did not show significant inaccuracy (Figure 1E). Although the mean inaccuracy of C and T bases was significant, the magnitude of this difference was less than 5% (Figure 1E-F). RNA editing is characterized by high variability in natural context (Harjanto et al. 2016). In addition, when RNA editing is induced by targeted RNA editing tools, in which RNA editing level is presumably quite similar between cells, there is variability in the reported editing of 5% or more (Cox et al. 2017; Vogel et al. 2018). In light of this, we consider our inaccuracy for C and T bases measurement marginal for most applications. Finally, we also compared the reliability of MultiEditR vs. NGS (RNAseq). We found a high coefficient of determination across all pooled bases relative to the identity line (Figure 3B), but when analyzing the difference in observed from expected measurements there were different trends of inaccuracies. Specifically, it appears that measuring C and T edits appears precise, but not completely accurate, while measuring G and A appears accurate, but not precise. However, it is important to note potential sources of error, some of which may be mitigated by further development of the MultiEditR algorithm

A potential source of error is random sampling error due to our sample size, therefore the reliability of our MultiEditR to NGS comparison could be improved by increasing the sample size and amplicon diversity of edited sites. Similarly, it would be advantageous to perform targeted amplicon deep sequencing for high coverage of these reads for comparison, as the level of coverage seen in our NGS data may introduce error (Harjanto et al. 2016). Not only would this be informative for addressing the accuracy of EditR, but it would also provide insight to the RNA editing field as a whole regarding the accuracy of measuring percent editing from RNA-seq. Furthermore, the observed error appears to be base specific and especially reproducible in C and T edits (Figure 3C), therefore it is reasonable to assume that there is some predictable, underlying process conferring error. In this line of thought, initial analyses (data not shown) and previous reports (Carr et al. 2009) suggest there may be context specific dependencies of peak height that could be modelled to provide corrected measurements of editing. Ultimately though, it is important to emphasize that for most applications of MultiEditR an average error of 6.3% when measuring T bases is within an acceptable margin of error. Therefore, when using MultiEditR we recommend sequencing with a primer that makes the hypothetical edit measurable as C-to-T or T-to-C editing. Given the nature of increased type I error rates when performing multiple NHSTs (Noble 2009), we also assessed the false positive rate of MultiEditR. As expected, our analysis yielded substantial observed false positives (29.2%, Figure 3A) when the level of significance was set to α = 0.01. However, were able to achieve increased stringency of the statistical test by decreasing the input level of significance to 0.0001, which resulted in a false positive rate of 10% while maintaining a reasonable sample size. This suggests that accurate significance testing by MultiEditR is feasible with proper calibration of the input level of significance. To properly calibrate the NHST, future work will need to implement a multiple comparisons correction such as the Bonferroni (Salkind 2010) or Dunn-Sidak (Šidák 1967) method. Furthermore, future work will also need to assess the type II error rate, i.e. false negatives, and thus the statistical power of MultiEditR relative to edits confirmed by NGS.

The use of MultiEditR for detection and quantification of editing directly from standard fluorescence-based Sanger sequencing of amplicons leads to the possibility of validation of several editing sites in a short time and with low cost. In fact different editing sites can be robustly detected and reasonably precisely quantified in the same amplicon, with only one Sanger sequencing reaction. This is an important feature when using site-direct RNA editing approaches, where detection and quantification of multiple editing sites along the same transcript is essential to evaluate off-target activity (Vogel and Stafforst 2018). Of course, the number of amplicons to validate can be easily scaled up maintaining both speed and low cost.

MultiEditR is not exclusive for analysis of RNA editing. The possibility to define in the software the base change of interest (Figure 2B) allows the application of the tool to many other contexts, as for example methylation detection after bisulfite treatment in DNA, in which the methylated C can be detected as a T-to-C base change. Furthermore, this feature is particularly important considering a very recent observation that RNA modifications affect RNA polymerase and reverse transcriptase activity and fidelity (Potapov et al. 2018). RNA is capable of hosting a variety of chemically diverse modifications, which in contrast to RNA editing are not detectable by sequencing. This makes their analysis dependent on antibody pull down and other specific techniques. The observation by Potapov et al. that fidelity during retrotranscription can be altered specifically by a specific modification opens the possibility that RNA modification can be “read” by a specific reverse transcriptase and then detected by Sanger sequencing. Overall, we expect MultiEditR to become a very useful tool for the RNA editing community, initially, but also the RNA modification community more broadly.

## AVAILABILITY

MultiEditR web app will be available at https://moriaritylab.shinyapps.io/MultiEditR upon submission Code for running web app locally and recreating figures is available at https://github.com/MoriarityLab/MultiEditR

## AUTHOR CONTRIBUTIONS

M.K., R.P., and B.M. designed the experiments. M.B. and T.L. developed KO cell lines. R.P. performed titration experiments and sequencing. M.K. wrote the program. R.T. defined the pipeline for NGS data processing. M.K. and R.P. analyzed the data. M.K. and R.P. wrote the manuscript. R.P and B.M. supervised the research.

## ACKNOWLEDGEMENTS

We thank the Flow Cytometry unit of the Imaging and Cytometry Core Facility, German Cancer Research Center (DKFZ), for providing excellent sorting services. We also thank Nina Papavasiliou (DKFZ) and Derek Nedveck (Univeristy of Minnesota) for their support of this project (Particularly with Derek Nedveck’s help with questions surrounding R shiny). Finally we also thank Walker Lahr (University of Minnesota) for helpful conversations surrounding this topic.

