## Supplementary Figures and materials for "MultiEditR: An easy validation method for detecting and quantifying RNA editing from Sanger sequencing"

#### SUPPLEMENTARY MATERIALS

**Figure S1.** ADAR1 knock out validation:
Western blot of A549 clones and bulk, obtained after transduction with lenti-CRISPR-ADAR1 exon 2 (A) or lenti-CRISPR-ADAR1 exon 3 (B). In both cases A549 cells were stimulated with 200 U/ml of IFN-α (pbl assay science, cat. no.#11100-1) for 16 h. Lenti-NT is the non-editing transduction control and was generated by transducing cells with the same vector (LentiCRISPRv2) but containing a non-targeting sgRNA (see Table S1). β-Actin and Calnexin were detected using the following antibodies: - Rabbit Calnexin polyclonal antibody, Enzo Life science, cat. no.#ADI-SPA-865-F; - Mouse β-Actin monoclonal antibody, Sigma-Aldrich, cat. no.#A5441.

**Figure S2.** APOBEC1 knock out validation:
Sanger sequencing chromatograms for RAW 264.7 wild-type and APOBEC1 knock-out after RT-PCR amplification of B2m 3'UTR region (oligos #19-20). The complete absence of C-to-U editing in the knock-out along the sequenced region confirm the deficiency of APOBEC1 editing activity in our knock-out clone.

**Figure S3.** C-to-U editing in CmAG:
Sanger sequencing chromatograms from 293T cells transfected only with CmAG or with CmAG, APOBEC1 and RBM47. Amplification of Apob region from CmAG transcript (oligos #21-2) and following Sanger sequencing show abundant C-to-T editing in several sites along the sequenced section.

**pJET-CmAG-WT and pJET-CmAG-6x**

CTCCCACAACGAGGACTACACCATCGTGGAACAGTACGAACGCGCCGAGGGCCGCCA
CTCCACCGGCGGCATGGACGAGCTGTACAAGaagcttaAACAAGTAGCTGGTGCCAAGGA
AAAAATAACTTCTTTCATGGAAAATTATAGAATTACAGATAATGATGTACTAATTGCCATA
GATAGTGCCAAAATCAACTTCAATGAAAAA**C**TCTCTCAA**C**TTGAGACATACGCGATA**C**AA
TTTGATCAGTATATTAAAGATAATTATGATCCACATGA**C**TTAAAAAGAA**C**TATTGCTGAGA
TTATTGAT**C**GAATCATTGAAAAGTTAAAAATTCTTGATGAACAGTATCATATCCGTGTAAA
TCTAGCAAAATCAATCCATAATCTCTATTTATTTGTTGAAAACGTTGATCTTAACCAAGTC
AGTAGTAGTAACACCTCTTGGATCCAAAATGTGGATTCCAATTATCAAGTCAGAATCCAA
ATTCAAGAAAAACTACAGCAGCTCAGGACACAAATTCAGAATATAGACATTCAGCAGCTT
GCTGCAGAGGTAAAACGACAG

In **yellow** are highlighted the 6 Cs which are unedited (C) in pJET-CmAG-WT and edited (T) in the pJET-CmAG-6x.

**mCherry-mApob-eGFP (CmAG) coding region:**

ATGGTGAGCAAGGGCGAGGAGGATAACATGGCCATCATCAAGGAGTTCATGCGCTTCA
AGGTGCACATGGAGGGCTCCGTGAACGGCCACGAGTTCGAGATCGAGGGCGAGGGCG
AGGGCCGCCCCTACGAGGGCACCCAGACCGCCAAGCTGAAGGTGACCAAGGGTGGCC
CCCTGCCCTTCGCCTGGGACATCCTGTCCCCTCAGTTCATGTACGGCTCCAAGGCCTA
CGTGAAGCACCCCGCCGACATCCCCGACTACTTGAAGCTGTCCTTCCCCGAGGGCTTC
AAGTGGGAGCGCGTGATGAACTTCGAGGACGGCGGCGTGGTGACCGTGACCCAGGAC

TCCTCCCTGCAGGACGGCGAGTTCATCTACAAGGTGAAGCTGCGCGGCACCAACTTCC
CCTCCGACGGCCCCGTAATGCAGAAGAAGACCATGGGCTGGGAGGCCTCCTCCGAGC
GGATGTACCCCGAGGACGGCGCCCTGAAGGGCGAGATCAAGCAGAGGCTGAAGCTGA
AGGACGGCGGCCACTACGACGCTGAGGTCAAGACCACCTACAAGGCCAAGAAGCCCG
TGCAGCTGCCCGGCGCCTACAACGTCAACATCAAGTTGGACATCACCTCCCACAACGA
GGACTACACCATCGTGGAACAGTACGAACGCGCCGAGGGCCGCCACTCCACCGGCGG
CATGGACGAGCTGTACAAGaagcttaAACAAGTAGCTGGTGCCAAGGAAAAAATAACTTCT
TTCATGGAAAATTATAGAATTACAGATAATGATGTACTAATTGCCATAGATAGTGCCAAAA
TCAACTTCAATGAAAAACTCTCTCAACTTGAGACATACGCGATACAATTTGATCAGTATAT
TAAAGATAATTATGATCCACATGACTTAAAAAGAACTATTGCTGAGATTATTGATCGAATC
ATTGAAAAGTTAAAAATTCTTGATGAACAGTATCATATCCGTGTAAATCTAGCAAAATCAA
TCCATAATCTCTATTTATTTGTTGAAAACGTTGATCTTAACCAAGTCAGTAGTAGTAACAC
CTCTTGGATCCAAAATGTGGATTCCAATTATCAAGTCAGAATCCAAATTCAAGAAAAACT
ACAGCAGCTCAGGACACAAATTCAGAATATAGACATTCAGCAGCTTGCTGCAGAGGTAA
AACGACAGACCCGGGATCCACCGGTCGCCACCATGGTGAGCAAGGGCGAGGAGCTGT
TCACCGGGGTGGTGCCCATCCTGGTCGAGCTGGACGGCGACGTAAACGGCCACAAGT
TCAGCGTGTCCGGCGAGGGCGAGGGCGATGCCACCTACGGCAAGCTGACCCTGAAGT
TCATCTGCACCACCGGCAAGCTGCCCGTGCCCTGGCCCACCCTCGTGACCACCCTGAC
CTACGGCGTGCAGTGCTTCAGCCGCTACCCCGACCACATGAAGCAGCACGACTTCTTC
AAGTCCGCCATGCCCGAAGGCTACGTCCAGGAGCGCACCATCTTCTTCAAGGACGACG
GCAACTACAAGACCCGCGCCGAGGTGAAGTTCGAGGGCGACACCCTGGTGAACCGCA
TCGAGCTGAAGGGCATCGACTTCAAGGAGGACGGCAACATCCTGGGGCACAAGCTGG
AGTACAACTACAACAGCCACAACGTCTATATCATGGCCGACAAGCAGAAGAACGGCATC
AAGGTGAACTTCAAGATCCGCCACAACATCGAGGACGGCAGCGTGCAGCTCGCCGACC
ACTACCAGCAGAACACCCCCATCGGCGACGGCCCCGTGCTGCTGCCCGACAACCACTA
CCTGAGCACCCAGTCCGCCCTGAGCAAAGACCCCAACGAGAAGCGCGATCACATGGTC
CTGCTGGAGTTCGTGACCGCCGCCGGGATCACTCTCGGCATGGACGAGCTGTACAAGT
AA

In red mCherry, in yellow mouse Apob and in green eGFP.

**Genes analyzed with REDItools v1 for NGS comparison with MultiEditR**

B2m - mm10 chr2:122,147,686-122,153,083 (RAW264.7 data)
MAVS - hg19 chr20:3,827,446-3,856,770 (A549 data)
DDX58 (RIG-I) - hg19 chr9:32,455,300-32,502,734 (A549 data)
SSR3 - hg19 chr3:156,257,929-156,272,973 (A549 data)

### Supplementary Figures

### Figure S1

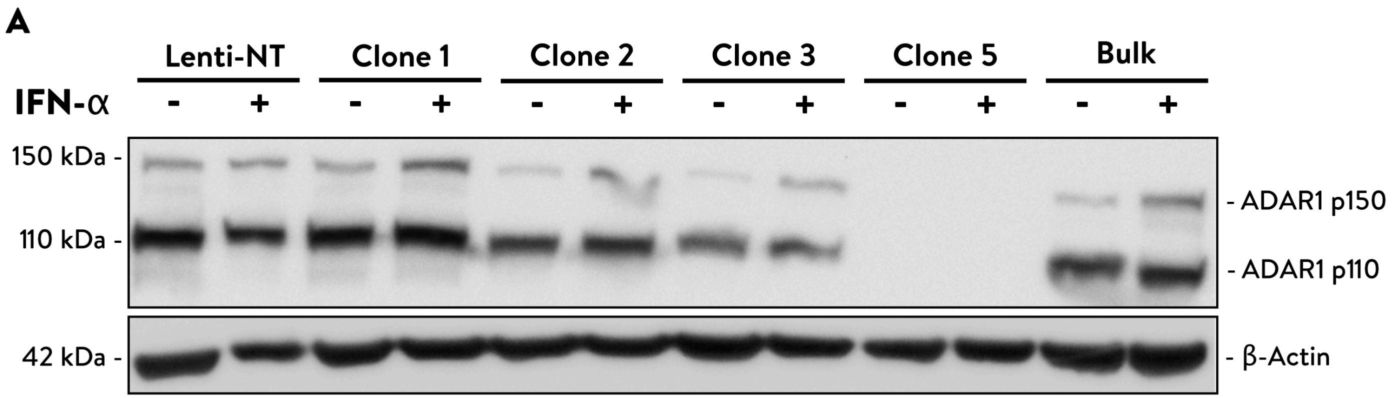
A
Lenti-NT
Clone 1
Clone 2
Clone 3
Clone 5
Bulk
IFN-α
- + - + - + - + - + - +
150 kDa -
110 kDa -
42 kDa -
- ADAR1 p150
- ADAR1 p110
- β-Actin

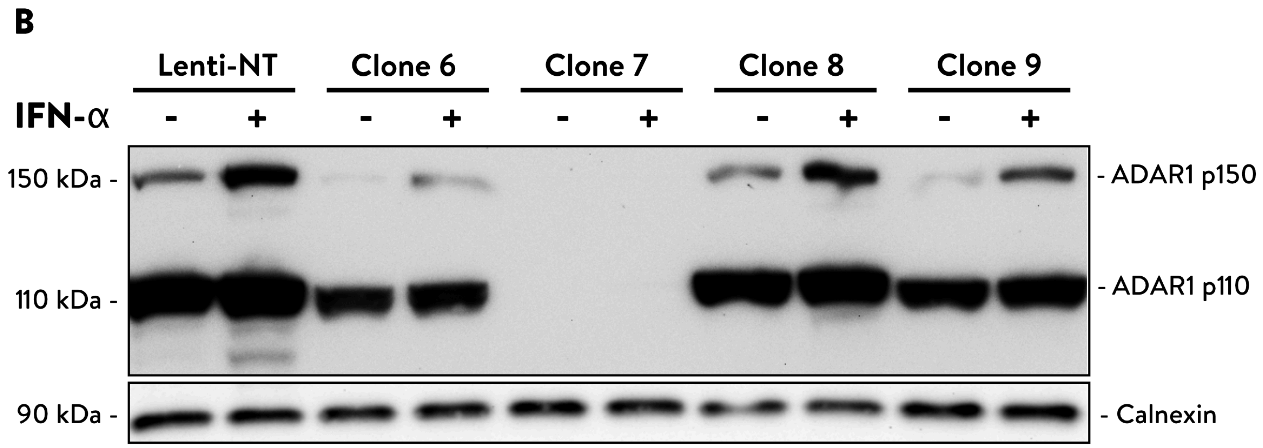
B
Lenti-NT
Clone 6
Clone 7
Clone 8
Clone 9
IFN-α
- + - + - + - + - +
150 kDa -
110 kDa -
90 kDa -
- ADAR1 p150
- ADAR1 p110
- Calnexin

Figure S2

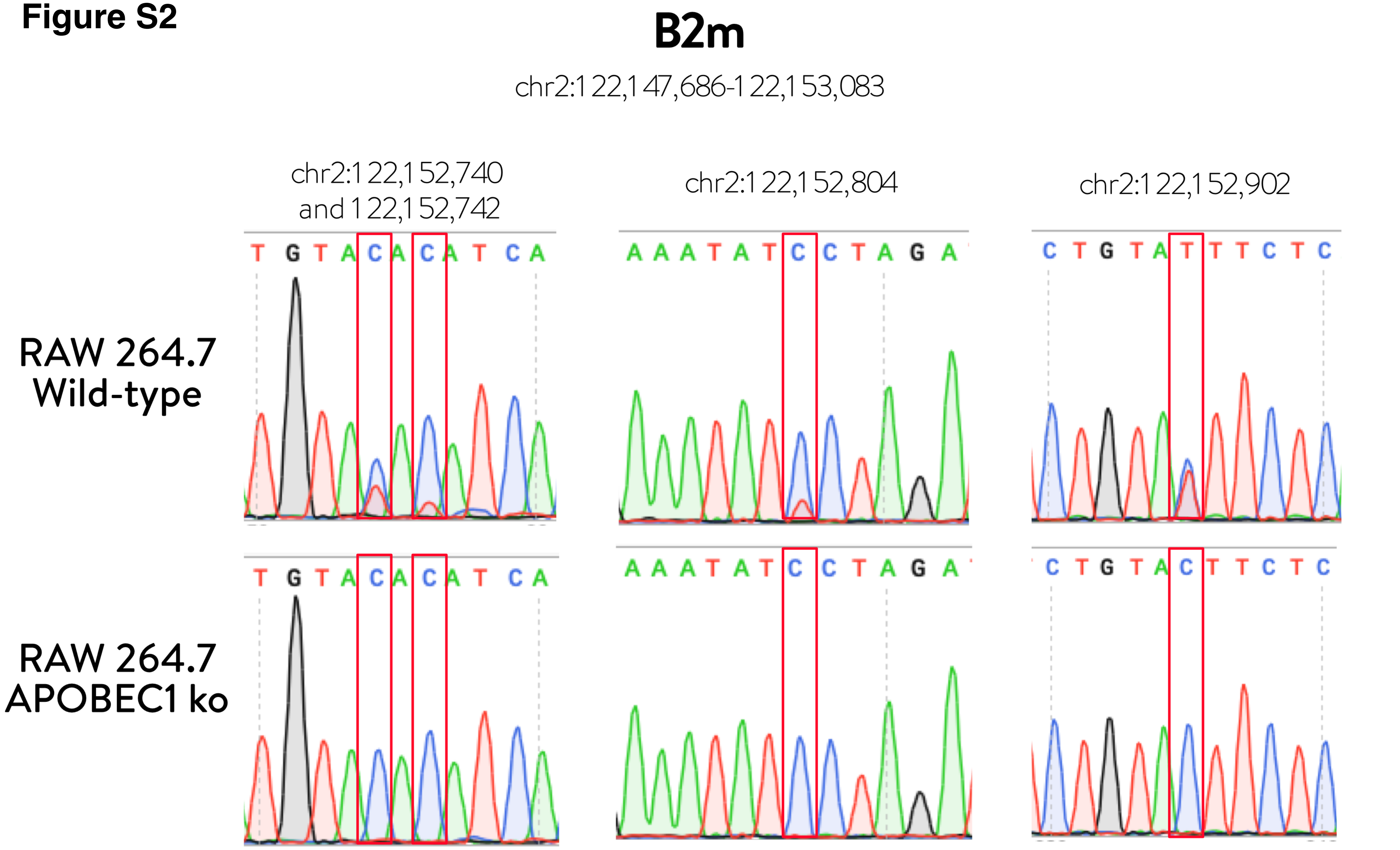
B2m
chr2:122,147,686-122,153,083
chr2:122,152,740
and 122,152,742
chr2:122,152,804
chr2:122,152,902
RAW 264.7
Wild-type
RAW 264.7
APOBEC1 ko
T G T A C A C A T C A
A A A T A T C C T A G A
C T G T A T T T C T C
T G T A C A C A T C A
A A A T A T C C T A G A
C T G T A C T T C T C

### Figure S3

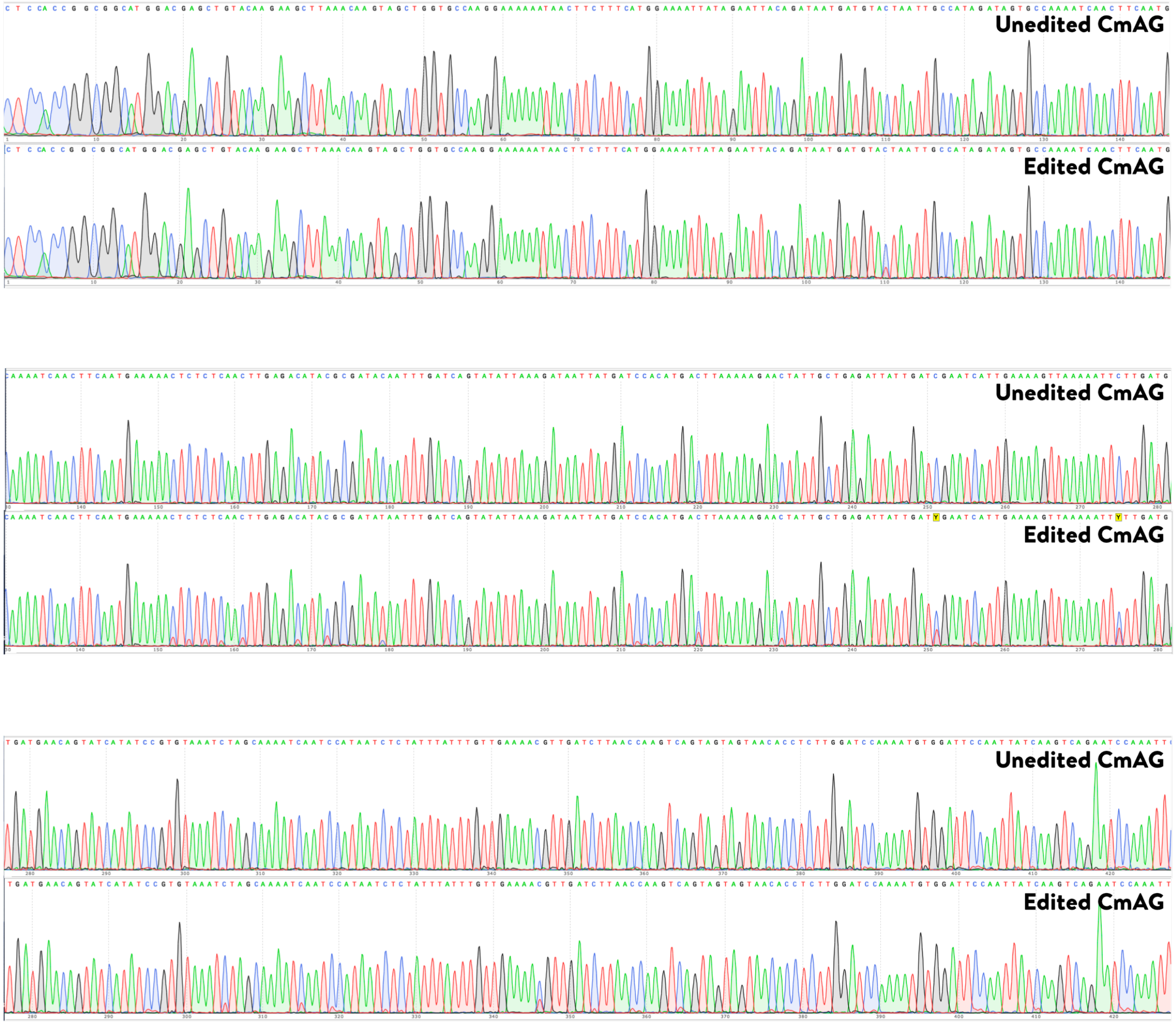
Unedited CmAG
Edited CmAG
Unedited CmAG
Edited CmAG
Unedited CmAG
Edited CmAG
